# Navigating in Clutter: How bumblebees optimize flight behaviour through experience

**DOI:** 10.1101/2025.03.13.643048

**Authors:** Manon Jeschke, Maximilian Stahlsmeier, Martin Egelhaaf, Olivier J. N. Bertrand

## Abstract

Bumblebees are excellent navigators that travel long distances while retracing paths to known locations. They forage not only in open terrains but also in cluttered environments where obstacles force them to deviate from direct paths. This study investigates the underexplored aspect of how bees become experienced foragers and optimize flight behaviour in cluttered terrains. We recorded flight trajectories of novice bees inexperienced in navigating cluttered environments and monitored their behavioural performance as they gained experience on subsequent foraging trips through numerous obstacles. By controlling for experience levels, we analysed how flight characteristics evolve with increasing expertise. Successful navigation in cluttered terrains requires avoiding collisions with obstacles. This is only possible if these can be detected by visual features such as the retinal displacement of contrast edges. Obstacles which are harder to detect and to avoid by the bees can affect their flight performance. By introducing transparent objects into our dense environment, we challenged collision avoidance and learning mechanisms, analysing their impact on flight optimization under different environmental conditions. Our findings reveal that experienced bees fly similar paths through clutter and quickly adapt their flight regardless of their training environment. However, the specific paths followed are influenced by environmental conditions. Transparent objects primarily affect naive bees’ flight patterns while having minimal impact on flight optimization, suggesting that the efficient flights of experienced bees result not solely from reflexive collision avoidance but from learning and previous experience in cluttered environments.

## Introduction

Navigating to a desired location is a crucial task in everyday life for animals across all taxa. Navigation takes place over a wide range of distances and can be accomplished by using different guiding systems (Buehlmann et al., 2020) and following different motivations, whether the animals are looking for a mating partner (Stepien et al., 2020), performing long-distance migration (Chowdhury et al., 2021) or finding back home (Gagliardo, 2013).

Bumblebees are skilful navigators and essential pollinators that play a crucial role in maintaining biodiversity and keeping the ecosystem functioning (Goulson et al., 2008; Potts et al., 2010). As central-place foragers, they regularly leave their nest to search for food sources and bring nectar and pollen back to their hive, requiring them to not only memorize nearby feeding sites, but also to retrace their path and find back to a previously visited location. During their foraging flights, these insects often navigate through complex, cluttered environments like a densely vegetated forests, as well as artificial environments such as tomato fields in a greenhouse, where the animal must deviate from a straight path to avoid collisions without drifting away from their overall goal direction (Crall et al., 2015). Efficient collision avoidance and navigation in these challenging environments are critical for the bumblebees’ foraging success, as collisions can cause injury or damage to their wings and therefore worsen their flight performance (Foster & Cartar, 2011; Mountcastle et al., 2016).

Previous studies have shown that insects, including bumblebees, rely on visual cues and optic flow for navigation and collision avoidance in complex environments (Baird et al., 2010; Chakravarthi et al., 2017; Lecoeur et al., 2019; Linander et al., 2015). In contrast to walking animals, which can use tactile information to detect obstacles, flying insects like bees rely on visual information from their eyes to detect and avoid obstacles (Ravi et al., 2019). The visual system of bumblebees is well-adapted to their foraging needs, with a relatively high spatial resolution in the frontal region of their eyes and the ability to perceive a wide range of colours (Dyer et al., 2008; Spaethe & Chittka, 2003).

Several studies have investigated the flight characteristics of experienced bumblebees in environments with various number of objects (i.e. various object density), showing that bees adapt their ground speed and axial velocity when they encounter obstacles (Burnett et al., 2020; Crall et al., 2015) and the variance of lateral excursion when bees fly through an obstacle field (Burnett et al., 2021). However, navigating in complex environments includes more than just reflexive reactions to avoid collisions with obstacles, which are mostly independent of experience. Also learning and establishing a route memory are key factors for bumblebees to optimize their flight (Burnett et al., 2021; Gonsek et al., 2021). Nevertheless, these studies focused on the flight behaviour of bees experienced with their specific cluttered environment. Therefore, it is not easily possible to deduce whether the flight around an obstacle is the consequence of reflexive collision avoidance or of previous experience with the cluttered terrain. Thus, the consequences of increasing experience and how it interacts with collision avoidance in bumblebees to optimize their flight behaviour are still poorly understood. To gain deeper insight into how collision avoidance and learning mechanisms interact to help bumblebees (*Bombus terrestris*) optimize their flight in dense environments, we recorded their flight behaviour while carefully controlling for prior experience. We trained bees in two different environments containing dark and transparent objects to challenge collision avoidance and learning mechanisms and understand how bumblebees adapt to different visual environments with increasing experience.

To the best of our knowledge, our study is the first to investigate the effect of experience on the optimization of flight characteristics in bumblebees navigating cluttered environments. Since bees are able to optimize their journey when collecting food from several spatially distributed sources on a single foraging trip (e.g. traplining in bumblebees see (Lihoreau et al., 2010)), we hypothesize that bumblebees will also optimize their flight behaviour with increasing experience in cluttered environments. However, the presence of transparent objects in the environment may increase the time the bee needs to learn a route and optimize their flight, since these objects are less salient and therefore harder to detect for the bees. Our findings contribute to a better understanding of how bumblebees become efficient foragers and navigate in cluttered environments, while simultaneously highlighting the role of both collision avoidance and experience in studies of insect navigation.

## Methods

### Experimental setup

The experimental setup consists of two flight tunnels, facilitating the connection between a bumblebee hive and a foraging chamber (see Figure 1A). Bumblebees exiting the hive accessed the flight tunnel for population transit through transparent tubes measuring 2.5 cm in diameter, alongside small acrylic boxes with dimensions of 8 × 8 × 8 cm^3^. The flight tunnel’s dimensions were 200 × 50 × 30 cm^3^. Notably, the walls and floor of the flight tunnel were covered with a red and white 1/f noise pattern, as previously described by (Ravi et al., 2019), to provide the bees with a pattern characterized by a natural like spatial frequency spectrum. Transparent acrylic was utilized for the tunnel ceiling to facilitate the observation of bee behaviour.

**Figure 1:**
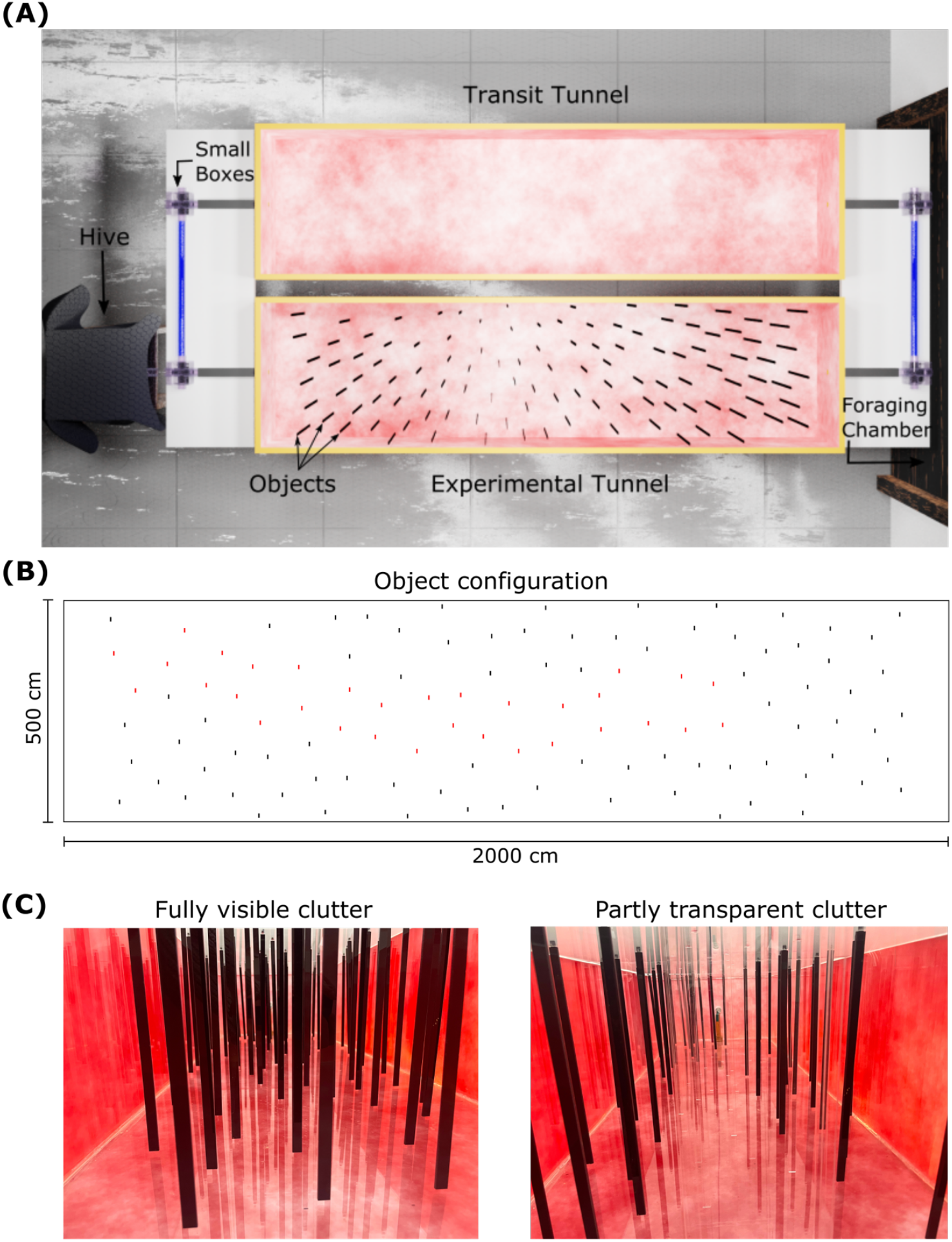
**(A)** Rendered top view of the setup. The hive was connected to the setup via small transparent boxes and tubes. Bees entered the foraging chamber (not shown) by flying through the transit tunnel (top). For the experiments, returning foragers were redirected into the experimental tunnel (bottom) where 110 vertical objects formed a complex cluttered environment the bees had to cross to return to the hive. **(B)** Object configuration. In the second environment, 32 objects (marked in red) were replaced with objects made from transparent acrylic. **(C)** Pictures taken from within the cluttered environment equipped with either all red objects (left) or transparent objects (right).

Upon their return from the foraging chamber, foragers were rerouted into the experimental tunnel for individual training. This tunnel was equipped with 110 randomly positioned objects, each measuring 29.5 × 1 × 0.3 cm^3^, thereby creating a cluttered environment. The objects had a minimal distance of 7 cm to each other to ensure that bees can pass the gaps without rotating along the yaw axis (Ravi et al., 2019). Notably, these objects were made from either transparent or red acrylic, with the latter designed to block light below 650 nm, as verified by spectrometer measurements (see Figure S1 in supplemental material), and therefore appearing dark to the bee (Dyer et al., 2008). The room and the flight tunnels were illuminated by light coming from the ceiling of the room. To capture the bee’s behaviour, four synchronized high-speed cameras (Basler acA2040-90umNIR) equipped with red filters (Heliopan RG715) were positioned above the tunnel at different angles, two top views and one side view on each end of the tunnel. To increase the visibility of the bees in the video, the tunnel was additionally illuminated from below using light filtered through a 650 nm cutoff low pass acrylic filter. This maximized the contrast on camera by creating a bright background while remaining natural lighting to the bees. Images were taken at a rate of 100 frames per second, allowing the bees to be detected, tracked, and their trajectories reconstructed in 3D.

### Animal handling

We used four colonies of *Bombus terrestris* from Koppert B.V. in The Netherlands. Upon receipt, the bees were transferred into a custom-designed acrylic box measuring 24×24×40cm^3^, which was draped with black cloth so that the nest is in the dark to imitate their natural environment. Within this enclosure, the bees were provided with pollen *ad libitum*, while having access to a 30% sucrose solution in a foraging area.

Foraging bees, i.e. bees undertaking flights between the hive and feeder were selected and individually marked. This process involved capturing the bee and immobilizing it within a marking tube. Subsequently, a numbered coloured plastic tag was fixed on the bee’s thorax using resin, allowing for individual identification. Upon tagging, the bee was released back into the experimental setup near the nest entrance.

### Experimental procedure

After a one-week habituation period, during which the bees had access to the foraging chamber by traversing the empty transit tunnel, we trained bees in two different cluttered environments and monitored their flight behaviour. On the outbound flight, bees flew through the empty transit tunnel into the foraging chamber. Bees returning from the foraging chamber were sequentially directed into the experimental tunnel, where we recorded their inbound flight through clutter until they reached the entrance to the hive. We trained two groups of bees in two different environments.

In the first environment, bees navigated through clutter composed solely of red acrylic objects, whereas in the second environment, 32 red objects were substituted with transparent ones (see Figure 1B, C). It is important to note that the locations of the objects and thus their density and arrangement were the same for both environments, with variations only in the visual clearance of certain sections of the clutter between environments.

In instances where a bee failed to locate the tunnel exit within five minutes, the recording was stopped, and the bee was returned to the hive. Each bee underwent 15 trials. Overall, we recorded the flight behaviours of 35 bees (16 in the first environment and 19 in the second environment), sourced from four distinct hives.

### Trajectories

Trajectories were captured using a custom written recording software developed in C++ (https://gitlab.ub.uni-bielefeld.de/neurobio-public/mcam-suite). The initial 100 frames were utilized to generate an image without the bee in the environment, i.e. a background image. Throughout the recording process, background subtraction was implemented, enabling the storage of cropped images, highlighting differences between the background and current frame, alongside their respective positions. Subsequently, we conducted image classification, retaining those images depicting bees. Furthermore, DeepLabCut (version 2.3.2) (Mathis et al., 2018) was employed to label the bee’s head and abdomen to determine the orientation of the bee based on the following dataset (https://pub.uni-bielefeld.de/record/2999609). We triangulated the bee’s 3D position by using synchronous detection of the bee on the images recorded by the different cameras, resulting in the 3D flight trajectory of the bee in the tunnel.

For analysis of flight characteristics, we excluded single trials where bees were walking through the clutter. For flight path similarity analysis, trajectory preparation involved removing the initial and final 25 cm to mitigate artefacts arising from the flight’s start and end. To analyse if experienced bees choose similar flight paths through clutter, we considered directed trajectories from the fifth trial onwards. Due to the choice of similarity measures (see below), we needed to exclude trajectories where bees flew back and forth in the tunnel.

## Data analysis and Statistics

To assess the similarity of individual flight paths across consecutive trials, we computed a similarity score utilizing distance functions. By comparing trajectories within each experimental group with each other using the Hausdorff distance, a distance matrix is generated. The Hausdorff distance is known as a geometry-based measure, representing the spatial similarity between two trajectories. A small distance value means that every point of either trajectory is close to some other trajectory points (Salarpour & Khotanlou, 2019). The similarity index represents the ratio of variance in distance values among trajectories of the same bee to the variance among randomly selected trajectories. A one sample Wilcoxon signed-rank test was used to determine whether the median similarity index differs from 1, indicating greater similarity among individual bee trajectories compared to randomly selected ones.

To explore to what extent bees optimize their flight behaviour through clutter with increasing experience, we analyse flight time, variance of flight speed, variance of lateral position, path sinuosity, average and variance of minimal distance to objects. These flight characteristics are highly relevant for bees, especially when flying through cluttered terrains. Using (generalized) linear mixed models (gmler package and r version 1.4.1) with trial number and experimental condition as fixed effects and bee ID as a random effect, we assess the impact of experience and environment on these flight characteristics. Furthermore, we assess differences in flight characteristics between groups before and after training using a Wilcoxon rank-sum test, with Bonferroni corrected p-values for multiple tests.

## Results

### Trajectories of bees differ between environments and level of experience

We uncover distinct behavioural patterns between naive and experienced bees as they navigate the cluttered environment (see Figure 2). During their initial trial, naive bees in both environments show exploratory behaviour, choosing indirect flight paths characterized by frequent back-and-forth movements within the tunnel. Surprisingly, we observe that collisions with objects are rare in both environments, regardless of whether part of the objects were transparent or not. Although the transparent objects are less salient and therefore should be harder to detect for the bees, they are proficient at steering clear of collisions. Naive bees tend to spend a considerable amount of time within the tunnel before returning to the hive. In contrast, experienced bees show a much more directed behaviour, displaying direct flight paths and efficiently crossing the cluttered area within a notably shorter time in their final training trial.

**Figure 2:**
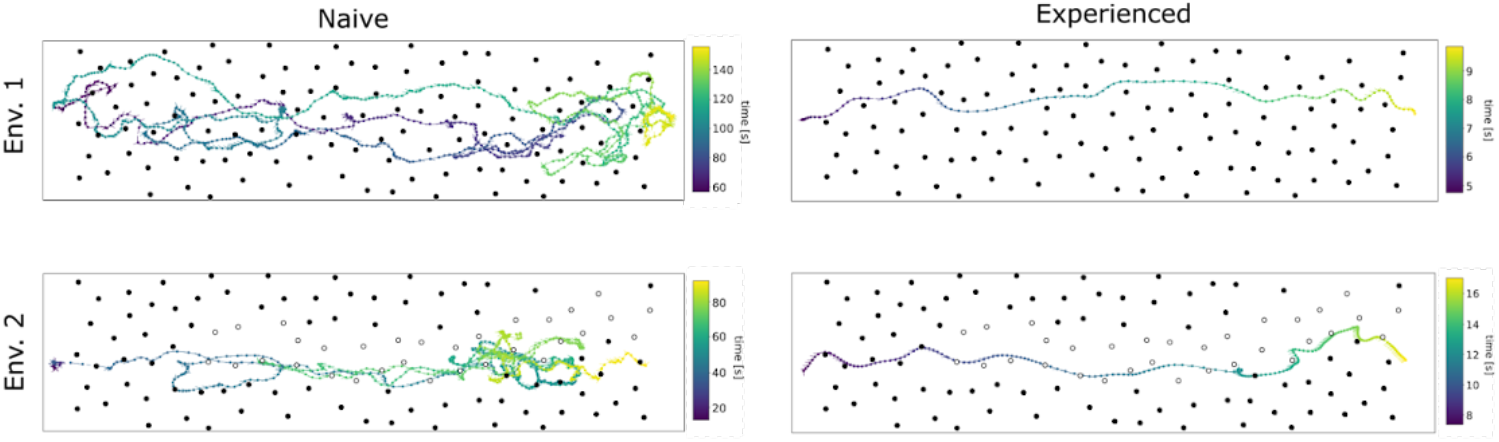
Example trajectories of naive and experienced bees in both environments. The bees are flying through the tunnel starting at the exit of the feeding chamber (left) to the entrance of the hive (right). The trajectories show the position and orientation of the bees. The time is color-coded. Objects are shown as circles. Empty circles refer to transparent objects.

After only a few training trials, bees show direct flights through both environments. However, we can observe that the paths taken differ if salient objects are exchanged along the tunnel by transparent ones. Trajectories of bees in environment 1 (see Figure 3A) occasionally cluster together to similar paths but are spread across a large part of the tunnel’s width while keeping distance to the walls. The mean lateral position of these trajectories is close to the midline of the tunnel but with a large variability (11.07 mm ± 62.26 mm). In contrast, trajectories in the environment with transparent objects (see Figure 3B) often converge and bees fly more frequently through the right half of the tunnel with a mean lateral position of -65.89 mm ± 31.92 mm.

**Figure 3:**
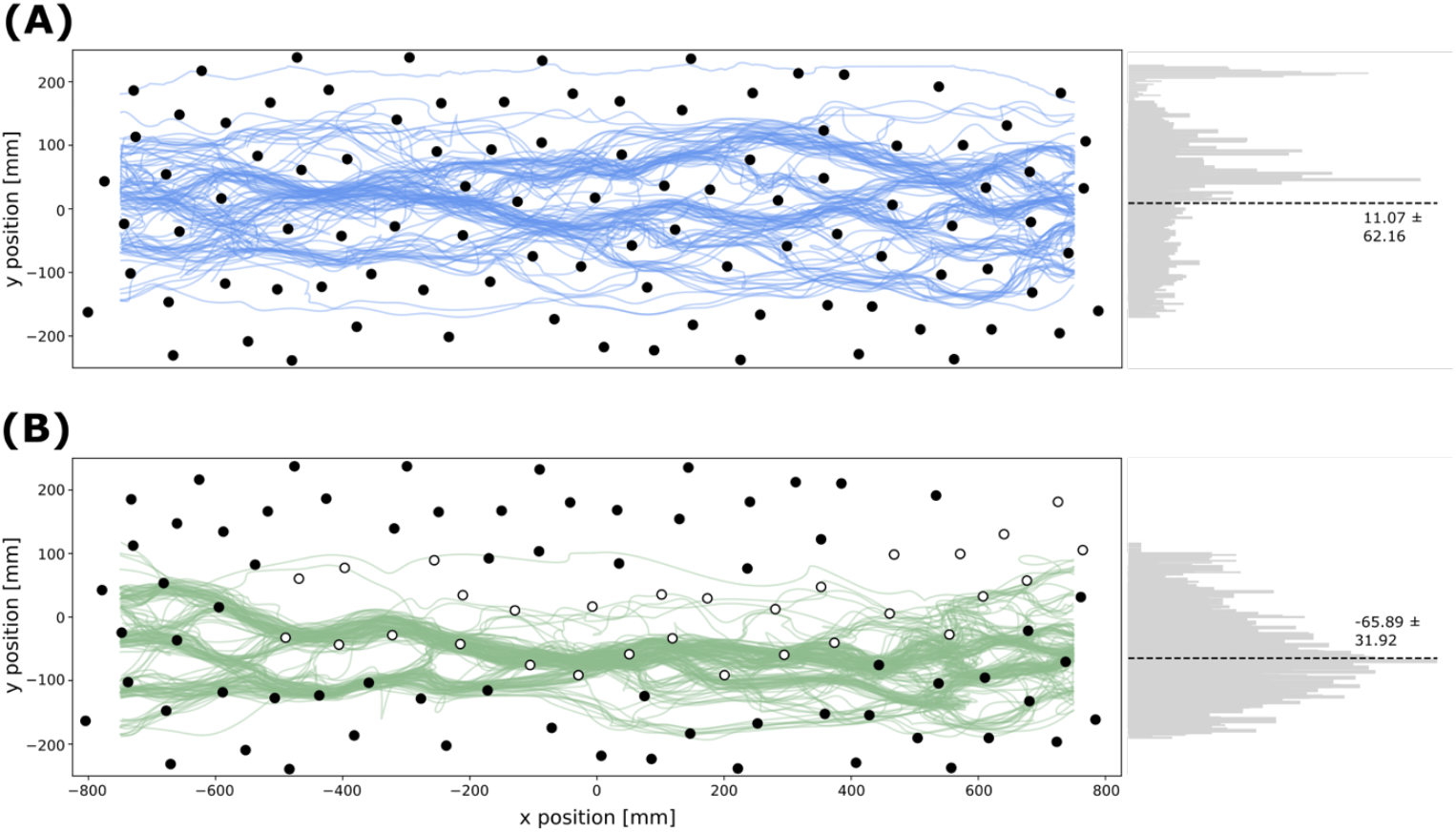
Directed trajectories from the 5th trial onwards of all bees in environment 1 (n = 105) (A) and environment 2 (n = 155) (B). The left subplot shows the trajectories of bees flying through the clutter. The object positions are indicated by filled and empty circles for dark and transparent objects respectively. The right subplot shows a histogram with the distribution of y positions. The dashed line indicates the average lateral position (mean ± std).

Comparing trajectories of individual bees of subsequent trials shows that they often fly along a similar path (see Figure 4A). The intra-individual difference of directed trajectories of the same bee is smaller than the inter-individual difference of trajectories across the whole population. On a population level, the similarity index which compares trajectories of the same bee with randomly chosen trajectories in the same environment is significantly smaller than 1 for both environments (environment 1: median = 0.83, p = 6.1035e-05; environment 2: median = 0.89, p = 1.9073e-06), indicating that the trajectories of individual bees are more similar than the trajectories across the whole population (see Figure 5B).

**Figure 4:**
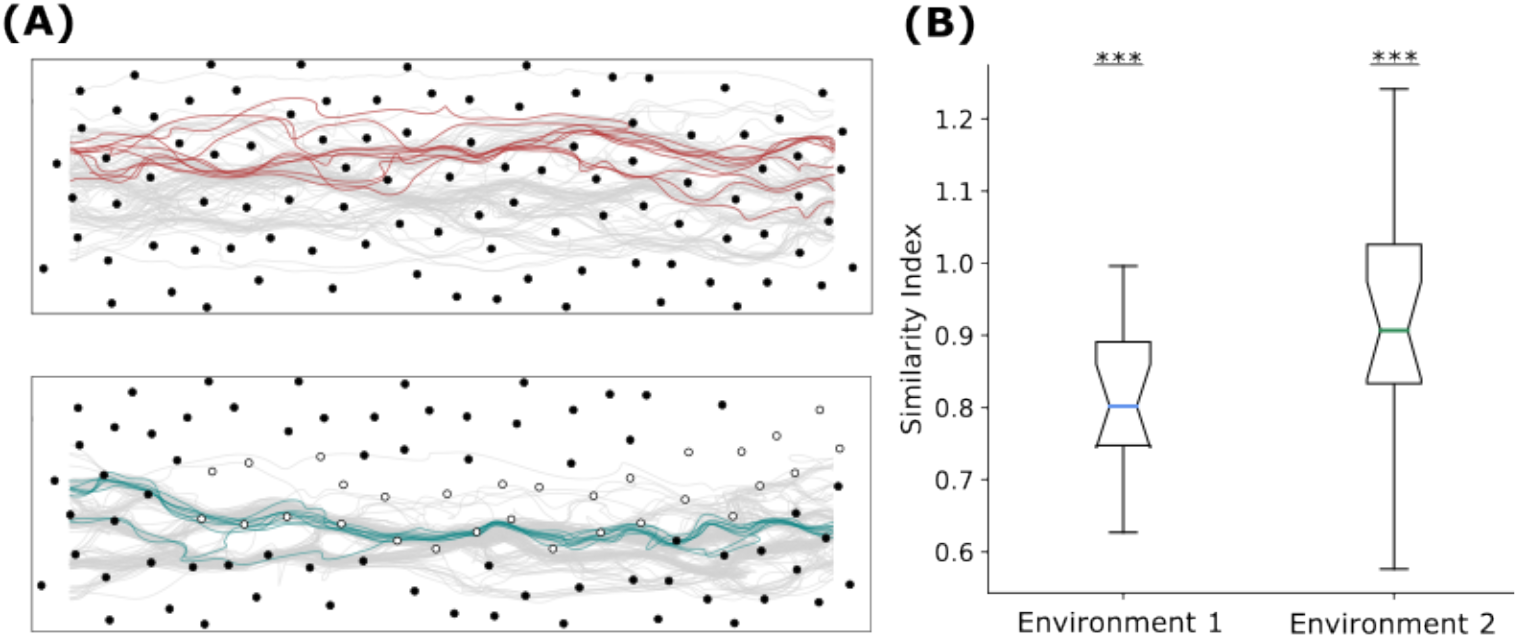
Similarity of flight paths of individual bees. (**A**) All directed trajectories from one example bee of environment 1 (top) and environment 2 (bottom) shown in colour. All directed trajectories of the respected group are shown in grey. Dark and transparent objects are represented as filled and empty circles. (**B**) Similarity index for both environments (n_Environment1_ = 16, N_Environment2_ = 19). The similarity index describes how similar trajectories of individual bees are compared to randomly selected trajectories of the population.

**Figure 5:**
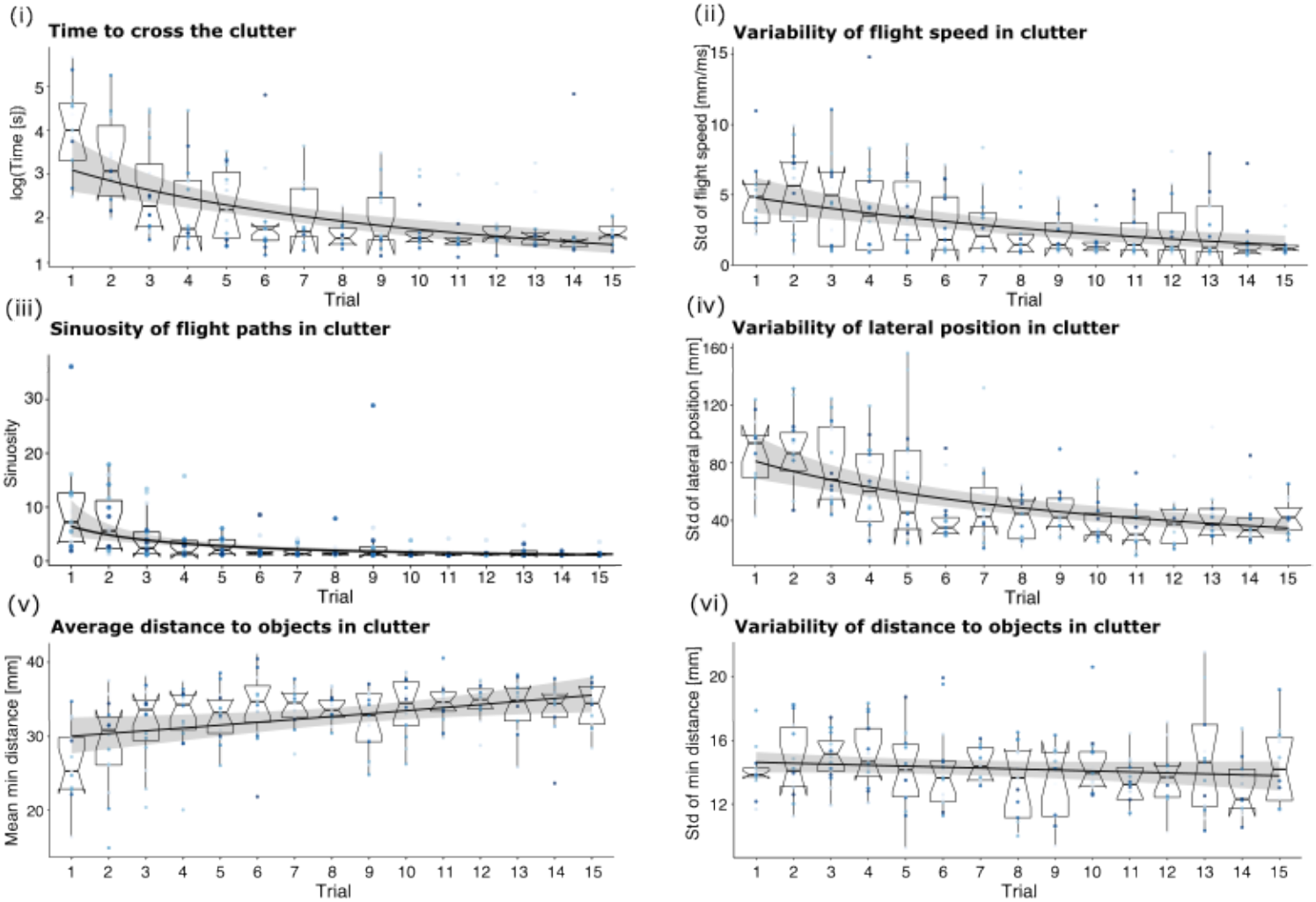
Effect of increasing experience on flight characteristics of bees in environment 1. Each subplot shows the development of one characteristic with increasing experience. Boxplots and individual-coloured points show measured data (n = 203), whereas the fit (black line) reflects the results of the GLMM performed for the statistical analysis and the 95% confidence interval (grey shaded area). For more details of the statistics see table 1 in supplemental material.

Overall, we conclude that bees quickly learn to efficiently cross different cluttered environments and travel along similar paths as they get more experienced in clutter.

### Bees optimize flight characteristics with increasing experience

We analyse the development of selected flight characteristics with increasing experience in both environments. Bees that were flying through the dense tunnel with only salient objects optimize most of the analysed flight characteristics with increasing experience (see Figure 5). They decreased the median time they needed to cross the tunnel over consecutive flights from 54 seconds to 5 seconds as well as the sinuosity of the flight paths. Moreover, their variability in flight speed decreases with increasing experience, indicating that more experienced bees vary their flight speed less during their flight and therefore perform less braking and acceleration manoeuvres while crossing the tunnel. The decrease in variability in the lateral position in the tunnel shows that bees use less the whole width of the tunnel while they are flying when they get more experienced with the clutter. The increase in the average minimal distance the bees keep to the objects during their flight suggests that they become more successful in avoiding obstacles. However, we observe no effect of experience on the variability in distance they keep to the objects in clutter.

Bees that were flying through a cluttered environment consisting of transparent and salient objects show similar trends in optimizing their flight with increasing experience in clutter (see Figure 6).

**Figure 6:**
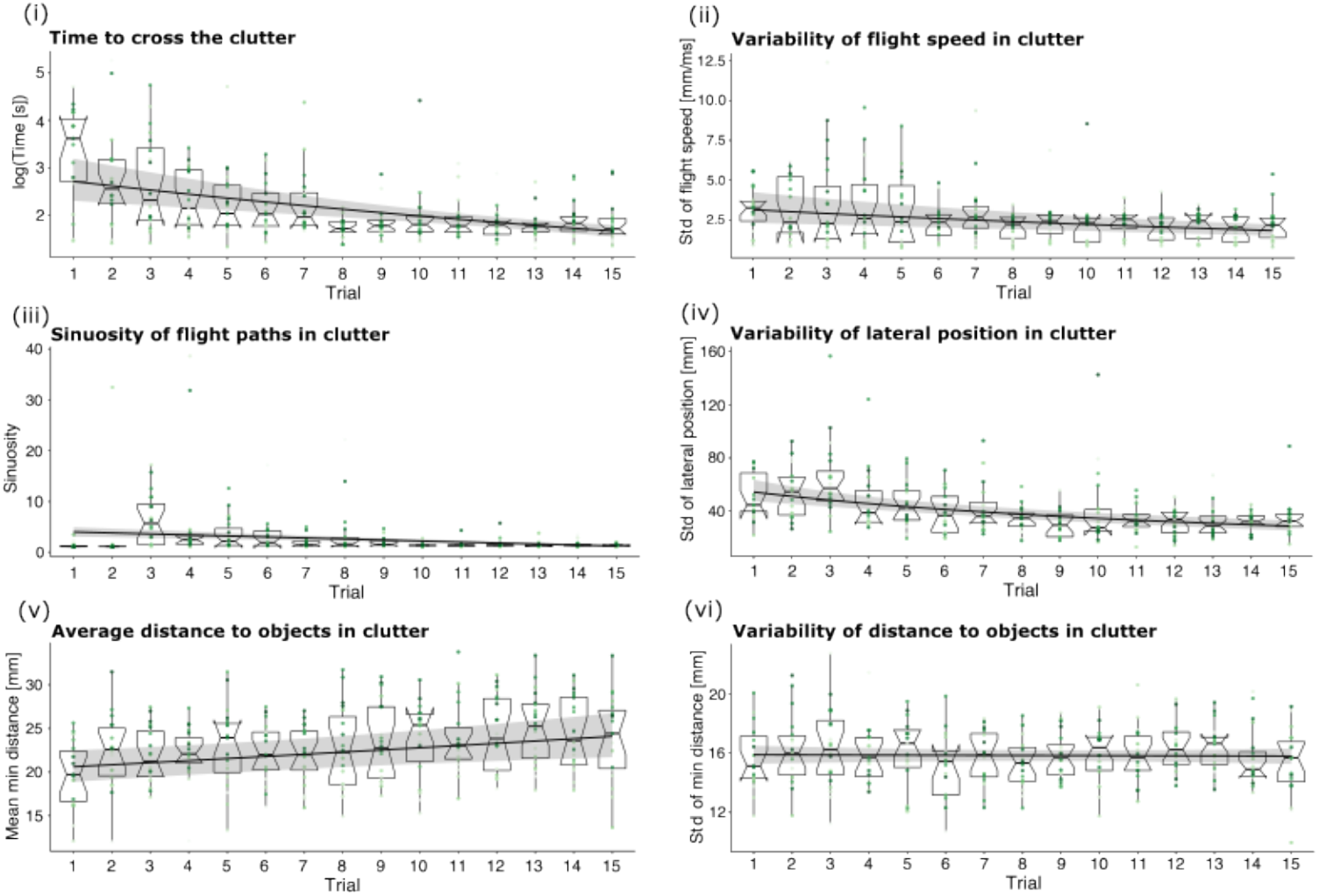
Effect of increasing experience on flight characteristics of bees in environment 2. Each subplot shows the development of one characteristic with increasing experience. Boxplots and individual-coloured points show measured data (n = 286), whereas the fit (black line) reflects the results of the GLMM performed for the statistical analysis and the 95% confidence interval (grey shaded area). For more details of the statistics see table 2 in supplemental material.

Although experience in clutter seems to be an important factor influencing the flight characteristics, it might not be the only one. Different bees might perform differently in clutter and could therefore lead to variation in the data. To test if individual bees that fly within the same environment optimize flight characteristics differently, we included the bee ID as a random effect in our GLMMs and compared model performances with and without the individual bees as a random effect. Notably, GLMMs that include the bee ID as a random effect perform better than models without a random effect, suggesting that individual bees differ in optimizing the investigated flight characteristics in both environments (see table 2 and 4 in supplemental material). This also corresponds to the finding that inter-individual variability between trajectories is larger than the intra-individual variability (see Figure 4).

To determine whether the level of experience or the dense environment or both is a significant predictor for the flight characteristics, we fit GLMMs with trial number and environment as fixed effects to the combined data of both environments. We fit models with either the trial number, the environment or both as a fixed effect and compare model performances using a likelihood-ratio-test. We find that models for all characteristics perform worse when we remove the trial as a fixed effect, indicating that the level of experience is a significant predictor for these flight characteristics (see table 12 in suppl. material for statistics). Moreover, we find that the environment is a significant predictor for the variance in lateral position, average minimal distance and variance in distance to objects, but not for the flight time, variance in flight speed and path sinuosity.

### Effect of the environment on novice bees

We found that the performance of the bees in terms of flying through a cluttered flight tunnel depends on the experimental conditions and in particular on whether some of the salient objects have been exchanged for transparent ones. This applies at least to some, though not all, of the flight characteristics analysed. Comparing the performance of bees in their first and last trial for all characteristics between environments reveals at which level of experience the environment has an effect on the flight behaviour.

The flight time and variability in flight speed show no difference between environments in naive and experienced bees (see Figure 7 A-B). Bees decrease flight time and variability in flight speed with increasing experience, but an environment with visually less dense areas does not impact naive or experienced bees in the time they need to cross the clutter nor the variability in flight speed.

**Figure 7:**
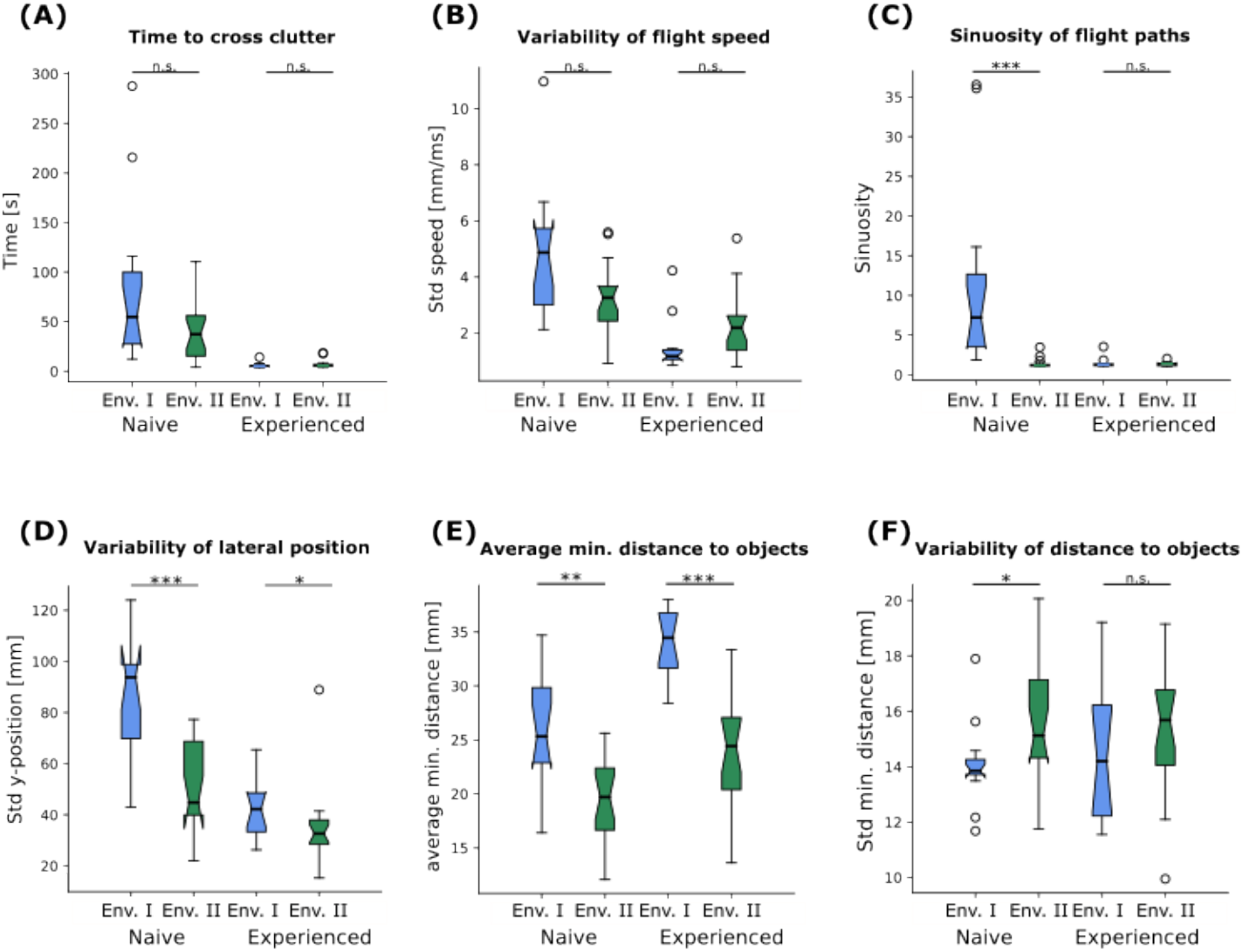
Comparison of flight characteristics for naive and experienced bees between environments. Panels A-F show different flight characteristics of naive bees on their first training trial (n_Environment1_ = 13, n_Environment2_ = 19) and experienced bees on their 15th trial (n_Environment1_ = 12, n_Environment2_ = 19). Within the same level of experience, we compare characteristics between environments using a Wilcoxon rank-sum test. P-values are corrected for multiple tests.

However, the environment influences how direct the bees cross the tunnel (see Figure 7 C-D). Naive bees in the partly transparent clutter show significantly straighter flight paths (Wilcoxon rank sum test: t = 4.5851, p = 4.5367e-06) than naive bees in the fully visible clutter. In addition, the variability in lateral position is significantly smaller for naive bees in the partly transparent clutter (Wilcoxon rank sum test: t = 3.5108, p = 0.0004) and also experienced bees vary their lateral position significantly less when they were trained in the partly transparent clutter (Wilcoxon rank sum test: t = 2.3522, p = 0.0186).

While these results might indicate that bees can cross a partly transparent clutter without problems of flying in between visually less salient obstacles, we observe that they nevertheless cope differently with salient and transparent objects (see Figure 7 E-F). Already naive bees keep significantly more distance to objects in the fully visible clutter than bees that fly through the partly transparent cluttered environment (Wilcoxon rank sum test: t = 3.4340, p = 0.0005). With increasing experience bees in both environments increase the average minimal distance they keep to objects in their environment. At the end of the training experienced bees in fully visible clutter keep on average more distance to objects than experienced bees in the partly transparent clutter (Wilcoxon rank sum test: t = 4.3394, p = 1.4285e-05). Interestingly, the variability in distance they keep to objects is smaller for naive bees in the fully visible clutter (Wilcoxon rank sum test: t = -2.4364, p = 0.0148), but is not different between environments for experienced bees.

In summary, our results revealed that environments with differently salient objects significantly influence bee flight behaviour. Notably, naive bees navigating our cluttered environment exhibited less straight flight paths, maintained closer proximity to objects, and showed reduced lateral position variation when encountering transparent objects.

## Discussion

By having bumblebees repeatedly fly through two different cluttered environments and challenging their collision avoidance abilities by confronting them with either only conspicuous objects or partially transparent objects, we were able to demonstrate their amazing navigational abilities in dense terrain. Our results show that bees rapidly learn to navigate successfully in an obstacle field and adapt their flight paths while progressively optimize flight characteristics. As they gain experience in cluttered spaces, bees in both environments fly along increasingly similar trajectories. Surprisingly, also bees that travelled through an environment containing also largely transparent objects managed to avoid collisions and optimized their flight with increasing experience, although the transparent objects in the environment provide less information for successful collision avoidance.

### Avoiding collisions with objects of different salience

Bees from both environments traversed the cluttered environment successfully. Contrary to initial expectations that bees might struggle to navigate through clutter containing transparent objects, they avoided collisions and maintained a safe distance from them.

The transparent objects in the environment might have had different brightnesses along their edges. These variations might have been detected by the bees. Several studies have shown that flying insects react to brightness changes induced by self-motion of the animal which create optic flow patterns on their retina (Egelhaaf, 2023; Li et al., 2017). Since closer objects create faster optic flow than more distant objects and an expanding optic flow pattern indicates an approaching obstacle, bees could use optic flow information in our experiment to estimate the relative distance to obstacles and perform avoidance manoeuvres. Even with transparent or low-contrast objects, this information could be used for collision avoidance, assuming that the bees detect brightness changes along the edges of the objects (Bertrand et al., 2015; Ravi et al., 2022). However, despite our behavioural findings that bees keep a certain distance to both visually conspicuous and transparent objects, the differences in minimal distance maintained between conspicuous and transparent objects suggest that bees still perceive transparent objects as less salient (see Figure 7 and suppl. Figure S2).

An analysis of individual trajectories reveals that bees not only learn to cross cluttered spaces on relatively direct routes but also tend to follow remarkably similar paths in subsequent trials. This intra-individual consistency highlights the bees’ ability to refine their navigation strategies through practice. The similarity of individual flight paths across trials aligns with recent studies on route stability in foraging bumblebees, a phenomenon known as traplining (Lihoreau et al., 2010), and similar route formation observed in natural foraging ants (Mangan & Webb, 2012).

The route formation we observe in our experiments might be a result of following a global direction combined with spontaneous collision avoidance manoeuvres, which is a rather reflexive behaviour and largely independent of experience, or route learning based on experience in the cluttered environment. These two aspects are not easy to distinguish experimentally. Therefore, we conducted experiments with non-transparent and transparent objects and challenged collision avoidance mechanisms and learning mechanisms to identify their relative role for bumblebees when optimizing their flight behaviour in cluttered terrains.

Previous modelling studies have shown that collision avoidance algorithms coupled with a goal direction can produce goal directed trajectories similar to those observed in navigating insects in clutter and the formation of a small number of routes through the simulated dense environment (Bertrand et al., 2015). Since collision avoidance is a reflexive behaviour and largely independent of experience, the bees in our experiment would only need to learn the goal direction, i.e. the exit of the tunnel. After an exploratory flight, we would expect that bees show direct flights through the cluttered environment in following trials. However, we rather observe that bees progressively optimize their flight and need multiple trials until they show direct flights through the clutter. Moreover, challenging collision avoidance mechanisms by using transparent objects in our environment leads to similar optimization of flight characteristics with increasing experience. Therefore, we conclude that collision avoidance mechanisms play a relatively small role in route formation in dense environments.

Since we observed the bees progressively optimized their flight with increasing experience in clutter, we assume that learning is involved in the process of flight optimization and route formation. Several papers have shown that insects can establish a spatial memory and strategically use learned visual landmarks for navigation (M. Collett et al., 2013). It is not entirely clear what visual features the bees learn and use for navigation in our paradigm, like panoramic views, single objects or object constellations as landmarks. This is not trivial given that the objects themselves do not represent useful landmark cues in our homogenous cluttered environment. In summary, we find few differences in the behaviour of bees between the different environments which suggests that collision avoidance plays a smaller relative role in flight optimization and route formation than the effect of experience in cluttered terrains.

### Optimization of flight behaviour in clutter

By carefully controlling for the bees’ level of experience in our cluttered environments, we could show that bumblebees can optimize their flight behaviour. This becomes evident by the reduction in time to cross the clutter, the decreased variability in flight speed, lower sinuosity of flight paths and distance to objects observed in both experimental groups.

While there has been little research about how flight characteristics evolve with increasing experience in cluttered terrains, tunnel experiments with experienced bees have been used a lot to study insect flight. Therefore, we can compare only our results of experienced bees to those of previous studies on flight characteristics of bumblebees.

In comparison with experiments using empty tunnels with a similar tunnel width as ours, our bees fly slower in the cluttered tunnels and show a larger variability in flight speed (Baird et al., 2010; Chakravarthi et al., 2017; Linander et al., 2015). Bees travel with a similar flight speed compared to ours when they fly through a tunnel with obstacles in their frontal field and have to perform lateral manoeuvres to avoid them (Burnett et al., 2020; Crall et al., 2015). However, they fly faster when the obstacles are in their lateral visual field and form a corridor the animals can fly through (Lecoeur et al., 2019). Therefore, we can strengthen the conclusion that flight speed in cluttered environments is highly influenced by the position of objects in the visual field.

We find numerous examples in the literature showing that bees that fly through an empty tunnel closely control their lateral position and fly along the midline with little variation in their lateral position (Chakravarthi et al., 2017; Linander et al., 2015, 2016). However, with objects positioned along the walls in the tunnel, the variability of lateral position decreases with increasing object density (Lecoeur et al., 2019) and bees increase precision with which they control their lateral position.

When our bees are trained to fly through a homogeneous cluttered environment of similar looking visually conspicuous objects, they fly on average in the centre of the tunnel but also show a large variation in their lateral position, likely due to the randomly placed objects they are confronted with in their lateral but also frontal visual field. Interestingly, a cluttered environment with transparent objects and, thus, visually less dense areas lead the bees to shift their average lateral position towards these areas and they show less variation in their lateral position.

The comparison of results of different experiments clearly shows that flight characteristics are influenced by environmental factors like the presence and placement of objects in the environment. Future experiments could investigate the potential influence of more environmental factors, such as the object density or spatial arrangement, on flight behaviour and help us to understand how flight characteristics evolve in different cluttered terrains.

### Different environments shape behaviour of naive bees

Our research reveals that flight behaviour is influenced not only by the animal’s experience but also by the environmental context. In contrast to naive bees in the environment with only visually conspicuous objects, naive bees flying through our second cluttered environment containing also transparent objects demonstrated more linear flight paths, reduced distance to objects, and less lateral position variation.

Comparing this behaviour to the existing literature, we find studies examining the behaviour of naive bees during their first outbound flights, also known as learning flights (Bertrand & Sonntag, 2023; T. S. Collett & Hempel de Ibarra, 2023). In contrast to the bees used in our experiments, that were only naive to our experimental tunnel containing the clutter, these bees were naive to the visual surrounding of their nest, but also to foraging, learning, navigating back to their nest etc., and can be described as fully naive bees. In a study examining bumblebee learning flights across different environmental densities, researchers found that fully naive bees, that were performing their first outbound flight, significantly alter their flight strategies in cluttered environments, prioritizing altitude gain over horizontal distance and showing more diverse orientation patterns, suggesting they adapt their flight altitude to acquire visual memories more effectively in complex landscapes (Sonntag, Lihoreau, et al., 2024). By varying the density of objects surrounding the nest entrance, they could show significant differences in flight behaviour. Since it has been shown that bumblebees do not only perform learning flights after they leave the nest for the first time but also perform similar flight manoeuvres when they find a new profitable food source (Robert et al., 2018). We might observe something similar when bees encounter a completely new environment, they are naive to, like our cluttered environment they encounter for the first time.

## Conclusion

In conclusion, our study reveals that bumblebees effectively optimize their flight behaviour in cluttered environments through experience. In addition to collision avoidance which constrains the way they traverse the clutter, they use their experience to adapt to the visual challenges posed by the clutter. Our findings highlight the ability of bees to refine the characteristics of their flight paths in response to environmental complexity. Notably, individual bees develop consistent trajectories over time, suggesting the formation of spatial memories and learned navigation strategies.

## Acknowledgements

We would like to thank Fabia Becker and Madelene Dombrowski for their help during the data collection.

## Notes

### Competing Interest Statement

The authors have declared no competing interest.

## References

Baird, E., Kornfeldt, T., & Dacke, M. (2010). Minimum viewing angle for visually guided ground speed control in bumblebees. The Journal of Experimental Biology, 213(Pt 10), 1625–1632.

Bertrand, O. J. N., Lindemann, J. P., & Egelhaaf, M. (2015). A Bio-inspired Collision Avoidance Model Based on Spatial Information Derived from Motion Detectors Leads to Common Routes. PLoS Computational Biology, 11(11), 1–28.

Bertrand, O. J. N., & Sonntag, A. (2023). The potential underlying mechanisms during learning flights. Journal of Comparative Physiology. A, Neuroethology, Sensory, Neural, and Behavioral Physiology, 209(4), 593–604.

Buehlmann, C., Mangan, M., & Graham, P. (2020). Multimodal interactions in insect navigation. Animal Cognition, 23(6), 1129–1141.

Burnett, N. P., Badger, M. A., & Combes, S. A. (2020). Wind and obstacle motion affect honeybee flight strategies in cluttered environments. The Journal of Experimental Biology. 10.1242/jeb.222471

Burnett, N. P., Badger, M. A., & Combes, S. A. (2021). Wind and route choice affect performance of bees flying above versus within a cluttered obstacle field. In bioRxiv (p. 2021.10.08.463704). 10.1101/2021.10.08.463704

Chakravarthi, A., Kelber, A., Baird, E., & Dacke, M. (2017). High contrast sensitivity for visually guided flight control in bumblebees. Journal of Comparative Physiology. A, Neuroethology, Sensory, Neural, and Behavioral Physiology, 203(12), 999–1006.

Chowdhury, S., Fuller, R. A., Dingle, H., Chapman, J. W., & Zalucki, M. P. (2021). Migration in butterflies: a global overview. Biological Reviews of the Cambridge Philosophical Society, 96(4), 1462–1483.

Collett, M., Chittka, L., & Collett, T. S. (2013). Spatial memory in insect navigation. Current Biology: CB, 23(17), R789–R800.

Collett, T. S., & Hempel de Ibarra, N. (2023). An “instinct for learning”: the learning flights and walks of bees, wasps and ants from the 1850s to now. The Journal of Experimental Biology, 226(6). 10.1242/jeb.245278

Crall, J. D., Ravi, S., Mountcastle, A. M., & Combes, S. A. (2015). Bumblebee flight performance in cluttered environments: Effects of obstacle orientation, body size and acceleration. The Journal of Experimental Biology, 218(17), 2728–2737.

Dyer, A. G., Spaethe, J., & Prack, S. (2008). Comparative psychophysics of bumblebee and honeybee colour discrimination and object detection. Journal of Comparative Physiology. A, Neuroethology, Sensory, Neural, and Behavioral Physiology, 194(7), 617–627.

Egelhaaf, M. (2023). Optic flow based spatial vision in insects. Journal of Comparative Physiology. A, Neuroethology, Sensory, Neural, and Behavioral Physiology. 10.1007/s00359-022-01610-w

Foster, D. J., & Cartar, R. V. (2011). What causes wing wear in foraging bumble bees? The Journal of Experimental Biology, 214(11), 1896–1901.

Gagliardo, A. (2013). Forty years of olfactory navigation in birds. The Journal of Experimental Biology, 216(Pt 12), 2165–2171.

Gonsek, A., Jeschke, M., Rönnau, S., & Bertrand, O. J. N. (2021). From Paths to Routes: A Method for Path Classification. Frontiers in Behavioral Neuroscience. 10.3389/fnbeh.2020.610560

Goulson, D., Lye, G. C., & Darvill, B. (2008). Decline and conservation of bumble bees. Annual Review of Entomology, 53, 191–208.

Lecoeur, J., Dacke, M., Floreano, D., & Baird, E. (2019). The role of optic flow pooling in insect flight control in cluttered environments. Scientific Reports, 9(1), 1–13.

Lihoreau, M., Chittka, L., & Raine, N. E. (2010). Travel optimization by foraging bumblebees through readjustments of traplines after discovery of new feeding locations. The American Naturalist, 176(6), 744–757.

Li, J., Lindemann, J., & Egelhaaf, M. (2017). Local motion adaptation enhances the representation of spatial structure at EMD arrays. PLoS Computational Biology, 13(12), 1–23.

Linander, N., Baird, E., & Dacke, M. (2016). Bumblebee flight performance in environments of different proximity. Journal of Comparative Physiology. A, Neuroethology, Sensory, Neural, and Behavioral Physiology, 202(2), 97–103.

Linander, N., Dacke, M., & Baird, E. (2015). Bumblebees measure optic flow for position and speed control flexibly within the frontal visual field. The Journal of Experimental Biology, 218(7), 1051–1059.

Mangan, M., & Webb, B. (2012). Spontaneous formation of multiple routes in individual desert ants (Cataglyphis velox). Behavioral Ecology: Official Journal of the International Society for Behavioral Ecology, 23(5), 944–954.

Mathis, A., Mamidanna, P., Cury, K. M., Abe, T., Murthy, V. N., Mathis, M. W., & Bethge, M. (2018). DeepLabCut: markerless pose estimation of user-defined body parts with deep learning. Nature Neuroscience, 21(9), 1281–1289.

Mountcastle, A. M., Alexander, T. M., Switzer, C. M., & Combes, S. A. (2016). Wing wear reduces bumblebee flight performance in a dynamic obstacle course. Biology Letters, 12(6). 10.1098/rsbl.2016.0294

Potts, S. G., Biesmeijer, J. C., Kremen, C., Neumann, P., Schweiger, O., & Kunin, W. E. (2010). Global pollinator declines: trends, impacts and drivers. Trends in Ecology & Evolution, 25(6), 345–353.

Ravi, S., Bertrand, O., Siesenop, T., Manz, L. S., Doussot, C., Fisher, A., & Egelhaaf, M. (2019). Gap perception in bumblebees. The Journal of Experimental Biology, 222(2), 1–10.

Ravi, S., Siesenop, T., Bertrand, O., Li, L., Doussot, C., Fisher, A., Warren, W. H., & Egelhaaf, M. (2022). Bumblebees display characteristics of active vision during robust obstacle avoidance flight. The Journal of Experimental Biology. 10.1242/jeb.243021

Robert, T., Frasnelli, E., Hempel de Ibarra, N., & Collett, T. S. (2018). Variations on a theme: bumblebee learning flights from the nest and from flowers. The Journal of Experimental Biology, 221(Pt 4). 10.1242/jeb.172601

Salarpour, A., & Khotanlou, H. (2019). Direction-based similarity measure to trajectory clustering. IET Signal Processing, 13(1), 70–76.

Sonntag, A., Lihoreau, M., Bertrand, O. J. N., & Egelhaaf, M. (2024). Bumblebees increase their learning flight altitude in dense environments. In bioRxiv. 10.1101/2024.10.14.618154

Sonntag, A., Sauzet, O., Lihoreau, M., Egelhaaf, M., & Bertrand, O. (2024). Switching perspective: Comparing ground-level and bird’s-eye views for bees navigating clutter. 10.7554/elife.99140

Spaethe, J., & Chittka, L. (2003). Interindividual variation of eye optics and single object resolution in bumblebees. The Journal of Experimental Biology, 206(Pt 19), 3447–3453.

Stepien, T. L., Zmurchok, C., Hengenius, J. B., Caja Rivera, R. M., D’Orsogna, M. R., & Lindsay, A. E. (2020). Moth Mating: Modeling Female Pheromone Calling and Male Navigational Strategies to Optimize Reproductive Success. NATO Advanced Science Institutes Series E: Applied Sciences, 10(18), 6543.

