## Supplemental figures and tables for "Navigating in Clutter: How bumblebees optimize flight behaviour through experience"

### Supplemental Material

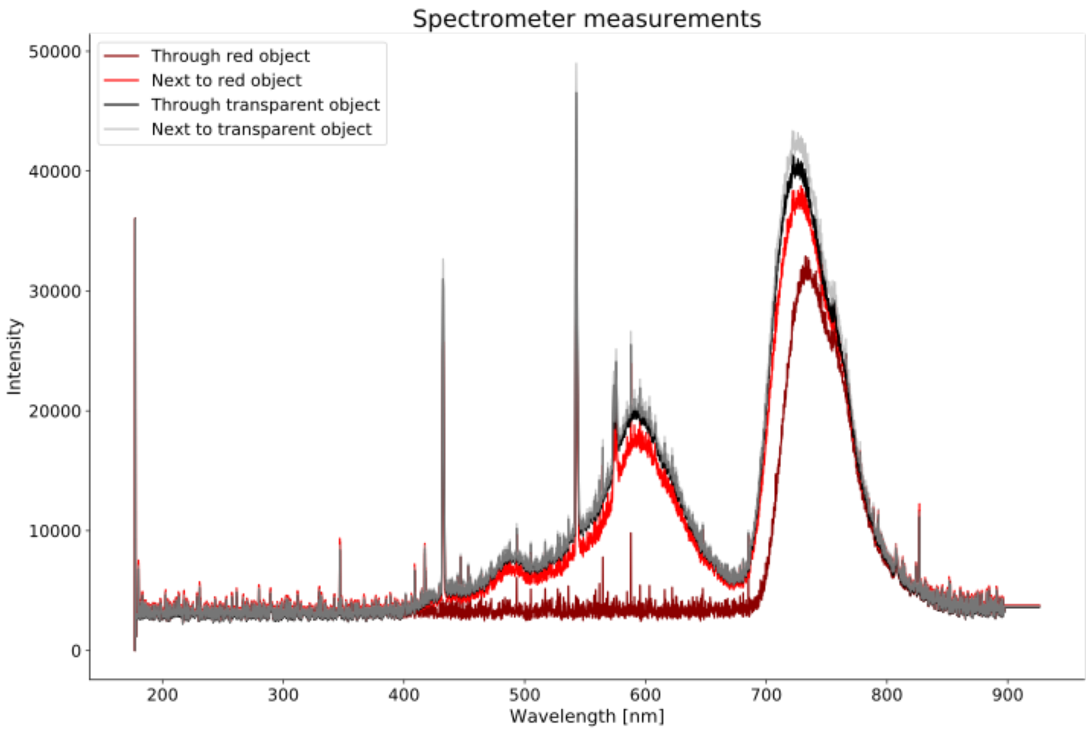


Suppl. Figure S1: Spectrometer measurements of different parts in the experimental setup. Using the same light conditions as during the experiments, we measured the light reflected by a dark red object (dark red curve), by a transparent object (black curve), the light inside the tunnel next to a dark red object (red curve) and the light inside the tunnel next to a transparent object (grey curve). Note that the red acrylic that was used to build the objects blocks light below 650nm. Therefore, the objects appear dark for the bees. The transparent objects allow all the light to pass through.

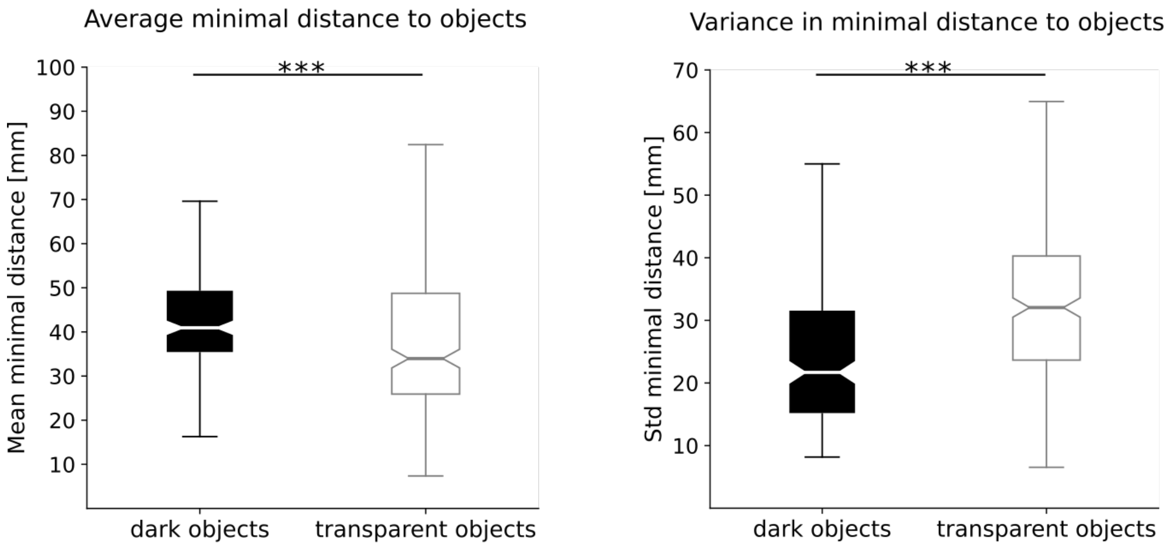


Suppl. Figure S2: Average and variability in minimal distance bees keep to different types of objects. In contrast to figure 6 and 7, we do not take all objects into account while calculating the minimal distance kept to object, but only the 32 objects that were replaced by transparent ones in the second environment.

Table 1: Mixed model results to explain the effect of experience on flight characteristics in the first environment

| | *Dependent variable:* | | | | | |
|---|---|---|---|---|---|---|
| | time | std speed | sinuosity | std y position | mean distance | std distance |
| | *generalized linear mixed-effects* | *generalized linear mixed-effects* | *generalized linear mixed-effects* | *generalized linear mixed-effects* | *generalized linear mixed-effects* | *linear mixed-effects* |
| | (1) | (2) | (3) | (4) | (5) | (6) |
| trial | $-0.052^{***}$ | $-0.086^{***}$ | $-0.274^{***}$ | $0.001^{***}$ | $0.012^{**}$ | $-0.061$ |
| | ($-0.075$, $-0.029$) | ($-0.123$, $-0.049$) | ($-0.372$, $-0.176$) | (0.001, 0.002) | (0.003, 0.021) | ($-0.149$, 0.028) |
| Constant | $1.129^{***}$ | $1.652^{***}$ | $5.153^{***}$ | $0.011^{***}$ | $3.390^{***}$ | $14.696^{***}$ |
| | (0.950, 1.307) | (1.366, 1.937) | (3.798, 6.507) | (0.009, 0.014) | (3.301, 3.478) | (13.991, 15.402) |
| Observations | 203 | 203 | 203 | 203 | 203 | 203 |
| Log Likelihood | $-192.340$ | $-373.778$ | $-387.076$ | $-892.761$ | $-584.477$ | $-443.158$ |
| Akaike Inf. Crit. | 396.680 | 759.556 | 786.153 | 1,797.521 | 1,180.954 | 898.315 |
| Bayesian Inf. Crit. | 416.560 | 779.435 | 806.032 | 1,817.401 | 1,200.833 | 918.195 |

*Note:* $^{*}p<0.1$; $^{**}p<0.05$; $^{***}p<0.01$

Table 2: Likelihood-ratio-test results to compare the goodness of fit of models including and excluding the Bee ID as a random factor in the first environment

<table>
<tr><th></th><th>#Df</th><th>LogLik</th><th>Df</th><th>Chisq</th><th>Pr(>Chisq)</th></tr>
<tr><td>log(time) ~ trial + (trial | beeid)</td><td>6</td><td>-192.34</td><td></td><td></td><td></td></tr>
<tr><td>log(time) ~ trial</td><td>3</td><td>-208.90</td><td>-3</td><td>33.12</td><td>3.040621e-07</td></tr>
<tr><td>speed_std ~ trial + (trial | beeid)</td><td>6</td><td>-373.78</td><td></td><td></td><td></td></tr>
<tr><td>speed_std ~ trial</td><td>3</td><td>-392.27</td><td>-3</td><td>36.99</td><td>4.614485e-08</td></tr>
<tr><td>sinuosity ~ trial + (trial | beeid)</td><td>6</td><td>-387.08</td><td></td><td></td><td></td></tr>
<tr><td>sinuosity ~ trial</td><td>3</td><td>-399.73</td><td>-3</td><td>25.31</td><td>1.331366e-05</td></tr>
<tr><td>y_std ~ trial + (trial | beeid)</td><td>6</td><td>-892.76</td><td></td><td></td><td></td></tr>
<tr><td>y_std ~ trial</td><td>3</td><td>-899.83</td><td>-3</td><td>14.14</td><td>0.0027</td></tr>
<tr><td>distobj_mean ~ trial + (trial | beeid)</td><td>6</td><td>-584.48</td><td></td><td></td><td></td></tr>
<tr><td>distobj_mean ~ trial</td><td>3</td><td>-596.98</td><td>-3</td><td>25.01</td><td>1.536575e-05</td></tr>
<tr><td>distobj_std ~ trial + (trial | beeid)</td><td>6</td><td>-443.16</td><td></td><td></td><td></td></tr>
<tr><td>distobj_std ~ trial</td><td>3</td><td>-446.78</td><td>-3</td><td>7.25</td><td>0.0643</td></tr>
</table>

Table 3: Mixed model results to explain the effect of experience on flight characteristics in the second environment

| | *Dependent variable:* | | | | | |
|---|---|---|---|---|---|---|
| | time | std speed | sinuosity | std y position | mean distance | std distance |
| | *generalized linear mixed-effects* | *generalized linear mixed-effects* | *generalized linear mixed-effects* | *generalized linear mixed-effects* | *generalized linear mixed-effects* | *linear mixed-effects* |
| | (1) | (2) | (3) | (4) | (5) | (6) |
| trial | $-0.035^{***}$ | $-0.038^{***}$ | $-0.198^{***}$ | $0.001^{***}$ | $0.286^{***}$ | $-0.007$ |
| | ($-0.048$, $-0.021$) | ($-0.055$, $-0.022$) | ($-0.272$, $-0.125$) | (0.001, 0.002) | (0.102, 0.469) | ($-0.068$, 0.054) |
| Constant | $1.034^{***}$ | $1.181^{***}$ | $4.209^{***}$ | $0.017^{***}$ | $20.756^{***}$ | $15.891^{***}$ |
| | (0.860, 1.208) | (0.866, 1.496) | (3.212, 5.206) | (0.014, 0.020) | (18.876, 22.637) | (15.237, 16.546) |
| Observations | 286 | 286 | 286 | 286 | 286 | 286 |
| Log Likelihood | −189.031 | −398.637 | −510.366 | −1,154.139 | −720.930 | −608.617 |
| Akaike Inf. Crit. | 390.062 | 809.275 | 1,032.733 | 2,320.278 | 1,453.861 | 1,229.234 |
| Bayesian Inf. Crit. | 411.998 | 831.211 | 1,054.669 | 2,342.214 | 1,475.796 | 1,251.170 |

*Note:* $^{*}p<0.1$; $^{**}p<0.05$; $^{***}p<0.01$

Table 4: Likelihood-ratio-test results to compare the goodness of fit of models including and excluding the Bee ID as a random factor in the second environment

| | #Df | LogLik | Df | Chisq | Pr(>Chisq) |
|---|---|---|---|---|---|
| log(time) ~ trial + (trial \| beeid) | 6 | -189.03 | | | |
| log(time) ~ trial | 3 | -232.30 | -3 | 86.54 | 1.213084e-18 |
| speed_std ~ trial + (trial \| beeid) | 6 | -398.64 | | | |
| speed_std ~ trial | 3 | -473.28 | -3 | 149.29 | 3.750611e-32 |
| sinuosity ~ trial + (trial \| beeid) | 6 | -510.37 | | | |
| sinuosity ~ trial | 3 | -524.48 | -3 | 28.23 | 3.247892e-06 |
| y_std ~ trial + (trial \| beeid) | 6 | -1154.14 | | | |
| y_std ~ trial | 3 | -1172.32 | -3 | 36.35 | 6.307203e-08 |
| distobj_mean ~ trial + (trial \| beeid) | 6 | -722.01 | | | |
| distobj_mean ~ trial | 3 | -808.72 | -3 | 173.41 | 2.341045e-37 |
| distobj_std ~ trial + (trial \| beeid) | 6 | -608.62 | | | |
| distobj_std ~ trial | 3 | -611.24 | -3 | 5.25 | 0.1541 |

Table 5: Mixed model results to explain the effect of experience on flight characteristics combined for both environments

| | *Dependent variable:* | | | | | |
|---|---|---|---|---|---|---|
| | time | std speed | sinuosity | std y position | mean distance | std distance |
| | *generalized linear mixed-effects* | *generalized linear mixed-effects* | *generalized linear mixed-effects* | *generalized linear mixed-effects* | *generalized linear mixed-effects* | *linear mixed-effects* |
| | (1) | (2) | (3) | (4) | (5) | (6) |
| trial | −0.041*** | −0.059*** | −0.232*** | 0.001*** | 0.012*** | −0.030 |
| | (−0.054, −0.029) | (−0.079, −0.039) | (−0.292, −0.172) | (0.001, 0.001) | (0.006, 0.017) | (−0.080, 0.021) |
| environment | 0.055 | −0.106 | 0.106 | 0.006*** | −0.376*** | 1.599*** |
| | (−0.069, 0.179) | (−0.440, 0.228) | (−0.231, 0.442) | (0.003, 0.008) | (−0.472, −0.281) | (1.004, 2.194) |
| Constant | 1.038*** | 1.441*** | 4.565*** | 0.011*** | 3.392*** | 14.471*** |
| | (0.892, 1.184) | (1.156, 1.726) | (3.718, 5.412) | (0.009, 0.013) | (3.309, 3.474) | (13.891, 15.051) |
| Observations | 489 | 489 | 489 | 489 | 489 | 489 |
| Log Likelihood | −395.951 | −799.701 | −898.367 | −2,047.660 | −1,309.299 | −1,051.636 |
| Akaike Inf. Crit. | 805.903 | 1,613.401 | 1,810.734 | 4,109.321 | 2,632.598 | 2,117.271 |
| Bayesian Inf. Crit. | 835.249 | 1,642.748 | 1,840.080 | 4,138.667 | 2,661.944 | 2,146.618 |

*Note:* *p<0.1; **p<0.05; ***p<0.01

Table 6: model comparison for flight time

| | *Dependent variable: time* | | |
|---|---|---|---|
| | (1) | (2) | (3) |
| trial | $-0.041^{***}$ | $-0.041^{***}$ | |
| | $(-0.054, -0.029)$ | $(-0.045, -0.038)$ | |
| environment | 0.055 | | 0.055 |
| | $(-0.069, 0.179)$ | | $(-0.068, 0.178)$ |
| Constant | $1.038^{***}$ | $1.070^{***}$ | $0.660^{***}$ |
| | (0.892, 1.184) | (1.064, 1.075) | (0.524, 0.797) |
| Observations | 489 | 489 | 489 |
| Log Likelihood | −395.951 | −396.324 | −407.551 |
| Akaike Inf. Crit. | 805.903 | 804.649 | 827.103 |
| Bayesian Inf. Crit. | 835.249 | 829.803 | 852.257 |
| *Note:* | | | $^{*}$p<0.1; $^{**}$p<0.05; $^{***}$p<0.01 |

Table 7: model comparison for variability in flight speed

| | *Dependent variable:Variability of flight speed* | | |
|---|---|---|---|
| | (1) | (2) | (3) |
| trial | $-0.059^{***}$ | $-0.059^{***}$ | |
| | $(-0.079, -0.039)$ | $(-0.079, -0.039)$ | |
| environment | −0.106 | | −0.115 |
| | $(-0.440, 0.228)$ | | $(-0.452, 0.222)$ |
| Constant | $1.441^{***}$ | $1.381^{***}$ | $1.003^{***}$ |
| | (1.156, 1.726) | (1.159, 1.603) | (0.652, 1.353) |
| Observations | 489 | 489 | 489 |
| Log Likelihood | −799.701 | −799.889 | −810.643 |
| Akaike Inf. Crit. | 1,613.401 | 1,611.777 | 1,633.286 |
| Bayesian Inf. Crit. | 1,642.748 | 1,636.931 | 1,658.441 |
| *Note:* | | | $^{*}$p<0.1; $^{**}$p<0.05; $^{***}$p<0.01 |

Table 8: model comparison for sinuosity

| | *Dependent variable: Sinuosity* | | |
|---|---|---|---|
| | (1) | (2) | (3) |
| trial | $-0.232^{***}$ | $-0.094^{***}$ | |
| | $(-0.292, -0.172)$ | $(-0.098, -0.091)$ | |
| environment | 0.106 | | 0.108 |
| | $(-0.231, 0.442)$ | | $(-0.262, 0.477)$ |
| Constant | $4.565^{***}$ | $1.606^{***}$ | $1.543^{***}$ |
| | (3.718, 5.412) | (1.603, 1.610) | (1.105, 1.980) |
| Observations | 489 | 489 | 489 |
| Log Likelihood | $-898.367$ | $-879.200$ | $-916.365$ |
| Akaike Inf. Crit. | 1,810.734 | 1,770.400 | 1,844.730 |
| Bayesian Inf. Crit. | 1,840.080 | 1,795.554 | 1,869.884 |
| *Note:* | | | $^{*}$p<0.1; $^{**}$p<0.05; $^{***}$p<0.01 |

Table 9: model comparison for variability in lateral position

| | *Dependent variable: Variability in lateral position* | | |
|---|---|---|---|
| | (1) | (2) | (3) |
| trial | $0.001^{***}$ | $0.001^{***}$ | |
| | (0.001, 0.001) | (0.001, 0.001) | |
| environment | $0.006^{***}$ | | $0.006^{***}$ |
| | (0.003, 0.008) | | (0.003, 0.009) |
| Constant | $0.011^{***}$ | $0.015^{***}$ | $0.018^{***}$ |
| | (0.009, 0.013) | (0.013, 0.017) | (0.014, 0.021) |
| Observations | 489 | 489 | 489 |
| Log Likelihood | $-2,047.660$ | $-2,053.178$ | $-2,064.523$ |
| Akaike Inf. Crit. | 4,109.321 | 4,118.357 | 4,141.045 |
| Bayesian Inf. Crit. | 4,138.667 | 4,143.511 | 4,166.200 |
| *Note:* | | | $^{*}$p<0.1; $^{**}$p<0.05; $^{***}$p<0.01 |

Table 10: model comparison for average distance to objects

| | *Dependent variable: Average distance to objects* | | |
|---|---|---|---|
| | (1) | (2) | (3) |
| trial | 0.012*** | 0.011*** | |
| | (0.006, 0.017) | (0.005, 0.017) | |
| environment | −0.376*** | | −0.378*** |
| | (−0.472, −0.281) | | (−0.474, −0.281) |
| Constant | 3.392*** | 3.173*** | 3.475*** |
| | (3.309, 3.474) | (3.070, 3.276) | (3.390, 3.559) |
| Observations | 489 | 489 | 489 |
| Log Likelihood | −1,309.299 | −1,323.601 | −1,314.992 |
| Akaike Inf. Crit. | 2,632.598 | 2,659.202 | 2,641.985 |
| Bayesian Inf. Crit. | 2,661.944 | 2,684.356 | 2,667.139 |
| *Note:* | | | *p<0.1; **p<0.05; ***p<0.01 |

Table 11: model comparison for variability in distance to objects

| | *Dependent variable: Variability in distance to objects* | | |
|---|---|---|---|
| | (1) | (2) | (3) |
| trial | −0.030 | −0.025 | |
| | (−0.080, 0.021) | (−0.076, 0.026) | |
| environment | 1.599*** | | 1.585*** |
| | (1.004, 2.194) | | (0.987, 2.183) |
| Constant | 14.471*** | 15.354*** | 14.258*** |
| | (13.891, 15.051) | (14.840, 15.869) | (13.805, 14.712) |
| Observations | 489 | 489 | 489 |
| Log Likelihood | −1,051.636 | −1,061.221 | −1,049.567 |
| Akaike Inf. Crit. | 2,117.271 | 2,134.442 | 2,111.133 |
| Bayesian Inf. Crit. | 2,146.618 | 2,159.597 | 2,136.287 |
| *Note:* | | | *p<0.1; **p<0.05; ***p<0.01 |

Table 12: Likelihood-ratio-test results to compare the goodness of fit of models including the trial number, the environment or both as fixed effects for combined data of both environments

<table>
<tr><th></th><th>#Df</th><th>LogLik</th><th>Df</th><th>Chisq</th><th>Pr(>Chisq)</th></tr>
<tr><td>(1) log(time) ~ trial + environment + (trial | beeid)</td><td>7</td><td>-395.95</td><td></td><td></td><td></td></tr>
<tr><td>(2) log(time) ~ trial + (trial | beeid)</td><td>6</td><td>-396.32</td><td>-1</td><td>0.746</td><td>0.387</td></tr>
<tr><td>(3) log(time) ~ environment + (trial | beeid)</td><td>6</td><td>-407.55</td><td>-1</td><td>23.199</td><td>1.46026e-06</td></tr>
<tr><td>(1) speedstd ~ trial + environment + (trial | beeid)</td><td>7</td><td>-799.70</td><td></td><td></td><td></td></tr>
<tr><td>(2) speedstd ~ trial + (trial | beeid)</td><td>6</td><td>-799.89</td><td>-1</td><td>0.375</td><td>0.539</td></tr>
<tr><td>(3) speedstd ~ environment + (trial | beeid)</td><td>6</td><td>-810.64</td><td>-1</td><td>21.89</td><td>2.894736e-06</td></tr>
<tr><td>(1) sinuosity ~ trial + environment + (trial | beeid)</td><td>7</td><td>-898.37</td><td></td><td></td><td></td></tr>
<tr><td>(2) sinuosity ~ trial + (trial | beeid)</td><td>6</td><td>-879.20</td><td>-1</td><td>74.33</td><td>0.0000</td></tr>
<tr><td>(3) sinuosity ~ environment + (trial | beeid)</td><td>6</td><td>-916.37</td><td>-1</td><td>36.00</td><td>1.976948e-09</td></tr>
<tr><td>(1) ystd ~ trial + environment + (trial | beeid)</td><td>7</td><td>-2047.66</td><td></td><td></td><td></td></tr>
<tr><td>(2) ystd ~ trial + (trial | beeid)</td><td>6</td><td>-2053.18</td><td>-1</td><td>11.036</td><td>0.0008</td></tr>
<tr><td>(3) ystd ~ environment + (trial | beeid)</td><td>6</td><td>-2064.52</td><td>-1</td><td>33.72</td><td>6.348872e-09</td></tr>
<tr><td>(1) distobj_mean ~ trial + environment + (trial | beeid)</td><td>7</td><td>-1309.30</td><td></td><td></td><td></td></tr>
<tr><td>(2) distobj_mean ~ trial + (trial | beeid)</td><td>6</td><td>-1323.60</td><td>-1</td><td>28.604</td><td>8.876971e-08</td></tr>
<tr><td>(3) distobj_mean ~ environment + (trial | beeid)</td><td>6</td><td>-1314.99</td><td>-1</td><td>11.39</td><td>0.0007</td></tr>
<tr><td>(1) distobj_std ~ trial + environment + (trial | beeid)</td><td>7</td><td>-1051.64</td><td></td><td></td><td></td></tr>
<tr><td>(2) distobj_std ~ trial + (trial | beeid)</td><td>6</td><td>-1061.22</td><td>-1</td><td>19.171</td><td>1.195148e-05</td></tr>
<tr><td>(3) distobj_std ~ environment + (trial | beeid)</td><td>6</td><td>-1049.57</td><td>-1</td><td>4.14</td><td>0.0419</td></tr>
</table>

Table 13: Wilcoxon rank-sum results comparing flight characteristics of bees between environments within the same level of experience

| | **level of experience** | **t-statistic** | **p-value** |
|---|---|---|---|
| time | naive | 1.8225 | 0.0683 |
| | experienced | -0.8922 | 0.3722 |
| std flight speed | naive | 2.0143 | 0.0439 |
| | experienced | -1.9466 | 0.0515 |
| sinuosity | naive | 4.5851 | 4.5367e-06 |
| | experienced | -0.2838 | 0.7764 |
| std lateral position | naive | 3.5108 | 0.0004 |
| | experienced | 2.3522 | 0.0186 |
| mean min. distance to objects | naive | 3.4340 | 0.0005 |
| | experienced | 4.3394 | 1.4285e-05 |
| std min. distance to objects | naive | -2.4364 | 0.0148 |
| | experienced | -1.0138 | 0.3106 |
